# The Human “Contaminome”: Bacterial, Viral, and Computational Contamination in Whole Genome Sequences from 1,000 Families

**DOI:** 10.1101/2022.01.31.478554

**Authors:** Brianna Chrisman, Chloe He, Jae-Yoon Jung, Nate Stockham, Kelley Paskov, Peter Washington, Dennis P. Wall

## Abstract

**Background:** The unmapped readspace of whole genome sequencing data tends to be large but is often ignored. We posit that it contains valuable signals of both human infection and contamination. Using unmapped and poorly aligned reads from whole genome sequences (WGS) of over 1,000 families and 5,000 individuals, we present insights into common viral, bacterial, and computational contamination that plague whole genome sequencing studies.

**Results:** We present several notable results: (1) In addition to known contaminants such as Epstein-Barr virus and phiX, sequences from whole blood and lymphocyte cell lines contain many other contaminants, likely originating from storage, prep, and sequencing pipelines. (2) Sequencing plate and biological sample source of a sample strongly influence contamination profile. And, (3) Y-chromosome fragments not on the human reference genome commonly mismap to bacterial reference genomes.

**Conclusion:** Both experiment-derived and computational contamination is prominent in next-generation sequencing data. Such contamination can compromise results from WGS as well as metagenomics studies, and standard protocols for identifying and removing contamination should be developed to ensure the fidelity of sequencing-based studies.

genomics WGS contamination

## Background

In the last decade, next-generation sequencing has become a commonly used tool in nearly every area of biology and has drastically changed the fields of human genomics [1, 2], metagenomics [3, 4], and pathogen surveillance [5, 6]. Additionally, open-source access to many bioinformatics tools [7, 8], benchmarking studies on the efficacy of computational pipelines [9, 10, 11], and improvements to laboratory procedures [12, 8] have made many next-generation sequencing use cases reliable or nearly reliable enough to be used clinically [13].

As the number of NGS studies grows as well as the diversity of sequencing sites and protocols, many studies have found various sources of contamination in publicly available NGS data. Such studies have found bacterial contamination in laboratory reagents and sequencing kits [14, 15], and common cross-contamination across samples [16]. In both genomic analysis of a single organism and metagenomics studies, such contamination can have critical impacts on downstream analysis. In whole-genome sequencing studies, bacterial contamination can result in false alignments and erroneous downstream variant calls [17, 18]. In studies of the microbiome, contamination can distort the estimation of microbial abundance of different genera [14, 19]. This is especially an issue for studies of microbiota that may have low microbial abundances, where even low levels of contamination may render metagenomics analysis inaccurate [20, 21].

In addition to improving laboratory protocols to reduce contamination [22, 23], several tools have been developed to identify contamination in next-generation sequencing data [24, 25, 26]. These tools rely on either sequencing a regent-only or blank sample to determine baseline contaminant levels of microbes, or rely on a measurement of total on-target DNA in a sample and assume an inverse relationship between on-target DNA biomass and contaminants. While such tools have improved the reliability of several microbiome studies, their assumptions can break down in sequencing experiments with many different confounding variables not included in the controls [27], very low abundance microbes [19], or when contaminate mass is comparable to sample mass [28]. In one particularly controversial microbiome, despite the numerous decontamination techniques available, researchers have been unable to agree on whether there is in fact a core blood microbiome or if all studies claiming so have simply been detecting contaminants [29, 30, 31, 32, 33].

More alarming than the presence of contamination in individual sequencing studies is the presence of contamination in reference databases. Studies have identified human DNA contamination in non-primate reference genomes [34], millions of contaminate sequences in GenBank [35], and human contamination in bacterial reference genomes that has created thousands of spurious protein sequences [36]. Such contamination risks compromising the findings of any genomics study, even if the researchers properly decontaminated or controlled for contaminants.

In order to better understand patterns of contamination in human whole genome sequencing, we analyzed sequences from the iHART dataset[37]. Originally curated to study genetic determinants of autism, the iHART dataset contains whole genome sequences from blood samples from children with autism, their siblings, and their parents, but also stands as an invaluable genomics resource due to its unique family structure [38, 39, 40]. iHART was sequenced at the New York Genome Sequencing Center, a common site for large sequencing studies, using commonly followed storage, prep, and sequencing protocols [37], making it a good model dataset to understand common sequencing issues. In addition to its unique family structure, the iHART collection contains both whole blood (WB) and lymphoblastoid cell lines (LCLs), and contains experimental batch information such as sequencing plate. By realigning reads from the iHART collection that were unmapped or poorly mapped to the human reference genome to a collection of viral, bacterial, and archael sequences, we are able to identify particular signatures of contamination that are unique to metadata variables.

We confirm the presence of many contaminating microbes that have been noted in other studies, including *Mycoplasma, Burkholderia, Bradyrhizobium, Mezorhizobium*, and *Variovorax*. We note that several microbes are strongly associated with cell type, suggesting that the LCL and WB storage pipelines may have differential effects on contamination signatures, and sequencing plate, suggesting that batch contamination can be a major risk to sequencing studies. Finally, we show that over 100 bacteria falsely associate with sex, indicating that reads from poorly catalogued regions of the sex chromosomes inaccurately map to bacterial contigs. We extract the offending k-mers that contribute to these mismappings, and suggest that researchers performing metagenomics pipelines on low microbial load environments filter their reads to remove such reads.

## Results

### Viruses and Bacteria Commonly Found In WGS

Following the basic pipeline shown in Fig. 1, Kraken2, a k-mer-based read classifier, classified many reads as belonging to bacteria and viruses (Fig. 2). The median number of reads per sample was 7.6×10^8^ [6.3×10^8^ - 8.9×10^8^ ]. Of the median 1.2×10^7^ [8.9×10^6^ - 1.7×10^7^] unmapped or poorly mapped reads per sample, a median of 37% [25%-44%] still matched to hg38 better than any other organisms, .03% [.01% - .08%] were reclassified as viruses, 21% [15% - 42%] as bacteria or archaea, 9% [2% - 17%] mapped ambiguously to organisms from multiple kingdoms, and 27% [19.2% - 40%] remained unmapped. Although some reads were classified ambiguously (with its set of k-mers matching equally well to sequences from multiple kingdoms), most reads were able to be classified to the species or strain level (58% [44% - 77%] of bacterial, viral, and archaeal reads that Kraken reclassified were reclassified to the species/strain level). Therefore, we aggregated our reads by their lowest taxonomic classification.

**Figure 1:**
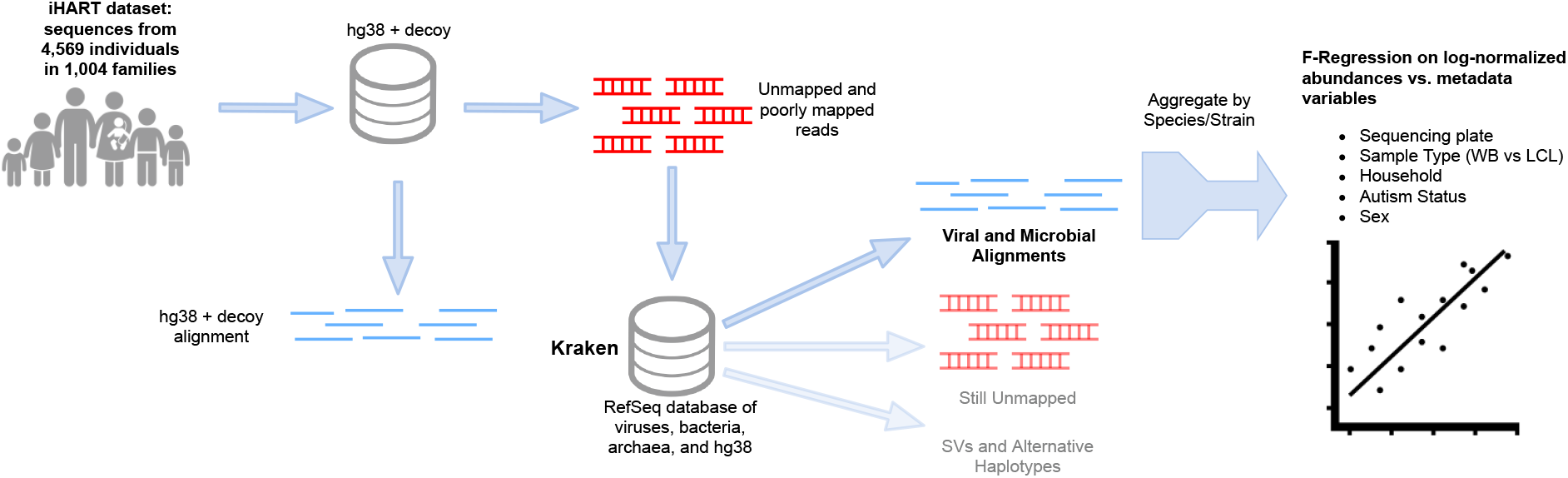
The general pipeline of the study: Reads from the iHART dataset that were unmapped or poorly aligned to hg38 were extracted and reclassified to a database of viruses, bacteria, archaea using Kraken. An F-regression was then performed on bacterial and viral counts against various sample-level metadata.

**Figure 2:**
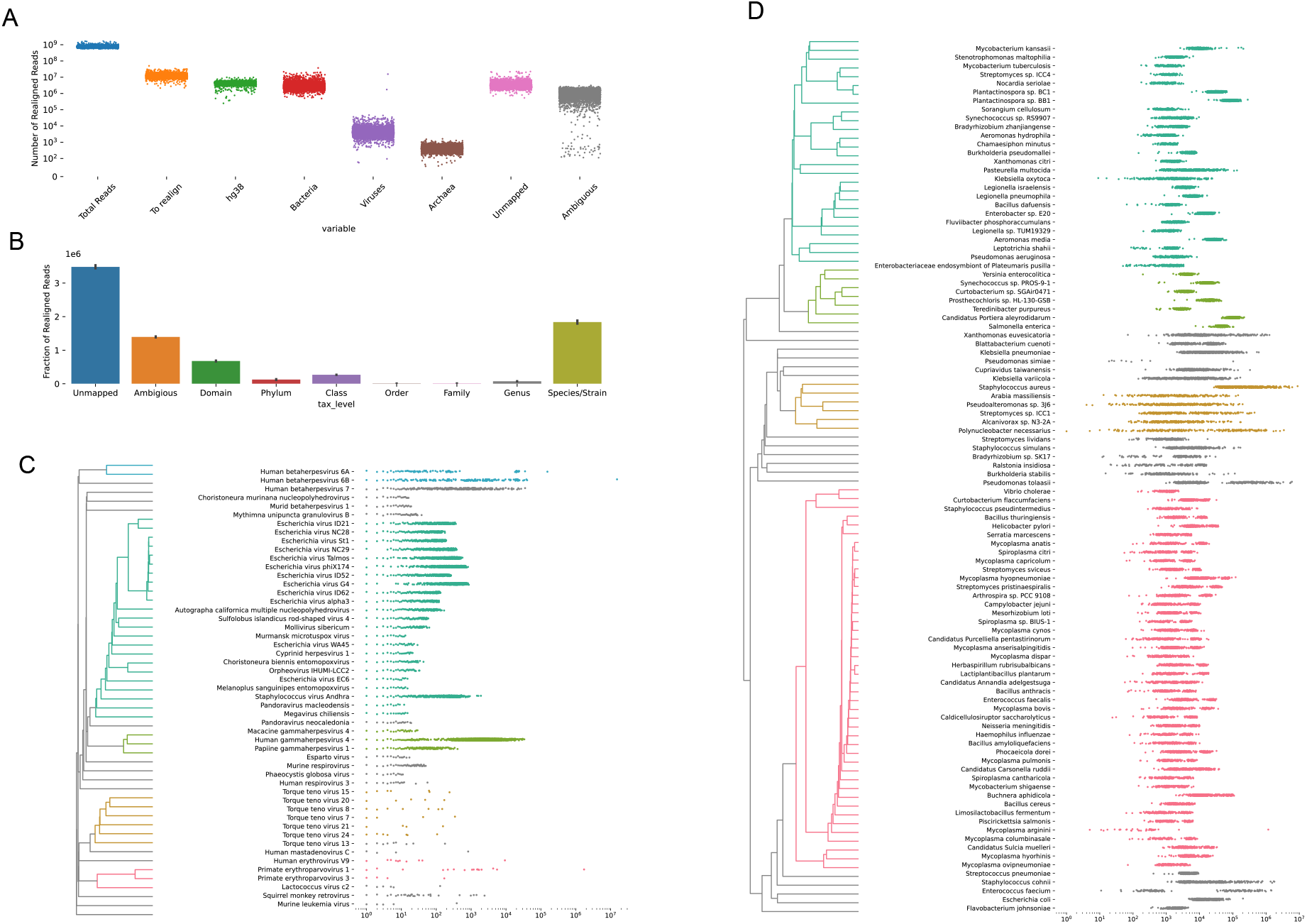
**(A)** Number of reads originally unmapped or poorly aligned to hg38 that Kraken2 classified as belonging to human, bacterial, viral, and archaeal sequences.**(B)** Taxonomic levels of Kraken2 classifications. While a significant fraction of reads were classified as ambiguous, Kraken2 was able to classify the majority of reads down to the species or strain level. **(C)** The top 50 most abundant viruses based on read count, clustered by Spearman correlation of sample abundances. Each point represents a sample’s read count. **(D)** Top 100 most abundant bacteria, archaea, and lower eukaryotes based on read counts and clustered by Spearman correlation of sample abundances.

We saw two main categories of viruses in the unmapped read space: DNA viruses likely originating from the human virome (such as human herpesviruses 6 and 7 as well as torque teno viruses), and common reagents used in the sequencing pipeline (such as lambda phage and herpesvirus 4). Phi X lambda phage is used as a spike-in for GC content in Illumina sequencing pipelines as well as to calibrate sequencing machines [41]. Herpesvirus 4, or Epstein Barr Virus (EBV) is used to immortalize LCLs [42]. Other phages or relatives to herpesvirus 4 are likely due to either mismappings, or commercial contamination, which we discuss more in the Discussion. Although the median number of reads belonging to viruses was small, our samples showed a wide range of viral read counts spanning over 4 orders of magnitude. The main contributors to this are lambda phage, which has a large variance across samples and EBV in which unsurprisingly LCL samples have much higher read counts over whole blood samples. Human herpesviruses 6A, 6B, and 7 also have large variances across samples, likely depending on whether an individual has a latent infection, active infection, or inherited chromosomally integrated human herpesvirus (iciHHV) [43].

We found many bacteria that were highly abundant in our samples. Notably, the top 100 most abundant bacteria also appeared in *>*90% of our samples, and most appeared in 100% of samples. These bacteria are almost certainly due to contamination, as even small traces of true bacteremia originally found in a blood sample would have been removed during sterilization steps in sample storage and prep, particularly in the LCL samples. In particular, we find many species of *Mycoplasmsa, Bradyrhizobium, Mycobacterium, Staphylococcus, Streptomyces, Streptococcus*, and *Pseudomonas* (Fig. 2C). Such bacteria are common water contaminants or either commonly found in human respiratory and oral cavities, and likely originated from reagent contamination or contamination introduced by a human experimenter. Many of the same contaminants we found were also found in other large scale WGS or metagenomics studies. We elaborate further on in the Discussion.

### Sample Type and Sequencing Plate Influence Microbial Contamination Profile

Using an forward F-regression, we found that sample type (LCL vs WB) and sequencing plate strongly influenced the abundances of many bacterial contaminants (Fig. 3). In particular, several species of *Achromobacter, Bradyrhizobium*, and *Burkholderia* were more abundant in whole blood samples (Fig. 3A), and several species of *Psuedomonas, Streptomyces*, and *Xanthomonas* were more abundant in LCL samples (Fig. 3B). Species of *Acidovorax, Bradyrhizobium, Mesorhizoium*, and *Variovorax* had different abundances according to sequencing plate (Fig. 3C).

**Figure 3:**
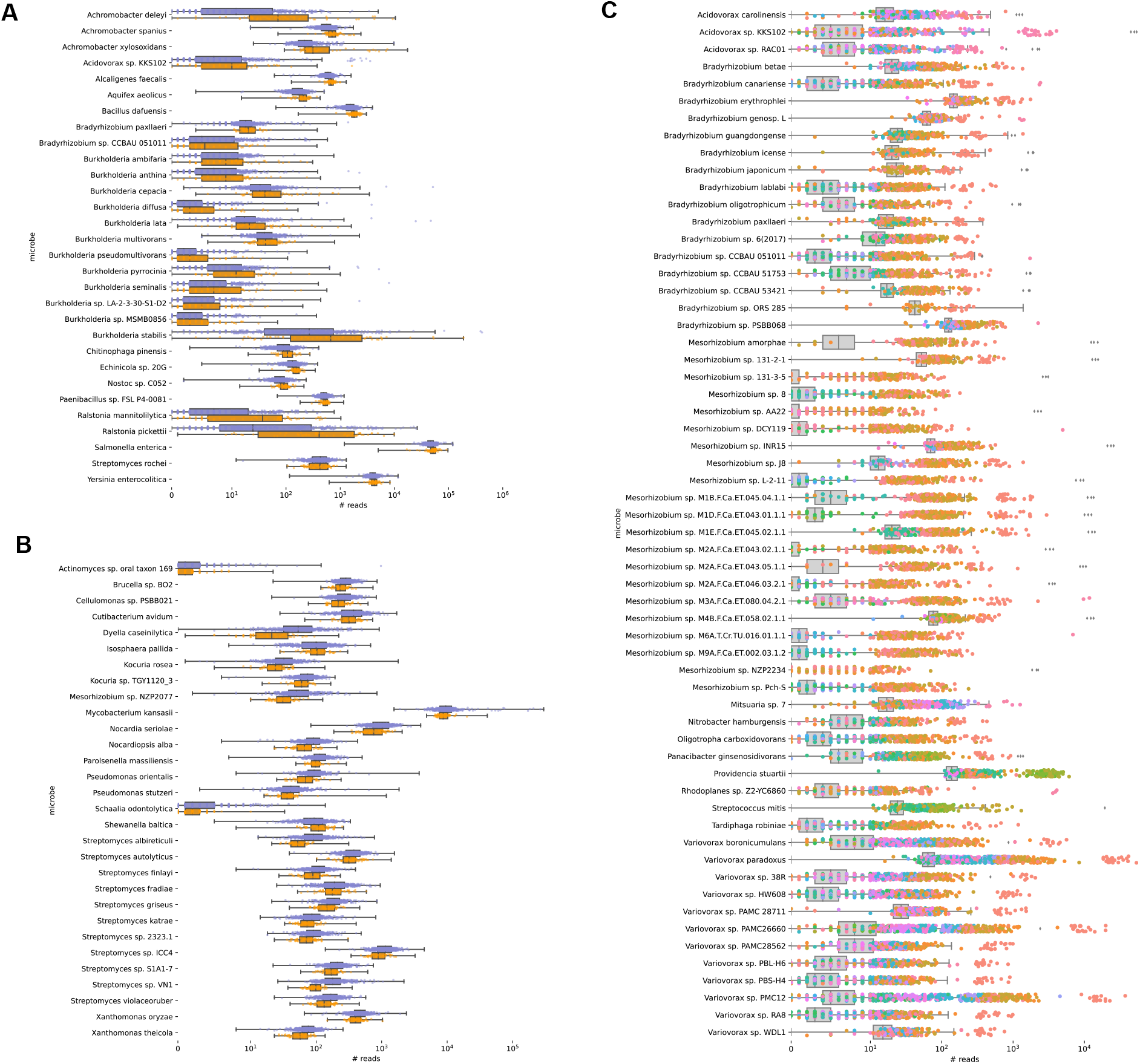
**(A)** Top 50 microbes most strongly associated with cell type per F-regression results, and more abundant in whole blood samples (orange) than LCL samples (purple). **(B)** Top 50 microbes most strongly associated with cell type per F-regression results, and more abundant in whole blood samples (orange) than LCL samples (purple). **(C)** Top 100 microbes most strongly associated with sequencing plate. Colors represent sequencing plates that had significantly higher abundances of a given microbe compared to the rest of the population. Samples from sequencing plates without significant enrichment of a microbe are captured in the grey box plots.

### Sex Chromosome Fragments Mismap to Bacterial Reference Genomes

Upon the F-regression showing many bacteria strongly associated with sex, we hypothesized that this was due to reads from the sex chromosomes being misclassified as bacteria. We show the male and female read counts for the bacteria most strongly associated with sex in (Figures 4A and 4B). Furthermore, we found that many bacteria had abundances strongly correlated between fathers and sons from the same nuclear family (an example is shown in 4D). The Y-chromosome has a notoriously poor reference genome with only about half its sequence present in hg38, and also has many repeats. We hypothesize that bacteria with high correlation between father and son read counts are due to repetitive regions in the Y-chromosome being misclassified as bacterial sequences. The number of repeats is passed down the male family line, and thus would be correlated between father and son. Interestingly, we also found that many bacteria had strong correlations between mother/daughter, and father/son (but not between father/daughter or mother/son) (Fig. 4C). An example of this is shown in Fig. 4 E). We hypothesize that reads mismapping to these bacteria may be coming from homologous sequences present on both X and Y chromosomes, with more repeats on the Y. Mother/daughter read counts would therefore show a mild correlation, but a father/daughter or mother/son correlation would get watered down by the large number of repeats coming from the Y-chromosome.

**Figure 4:**
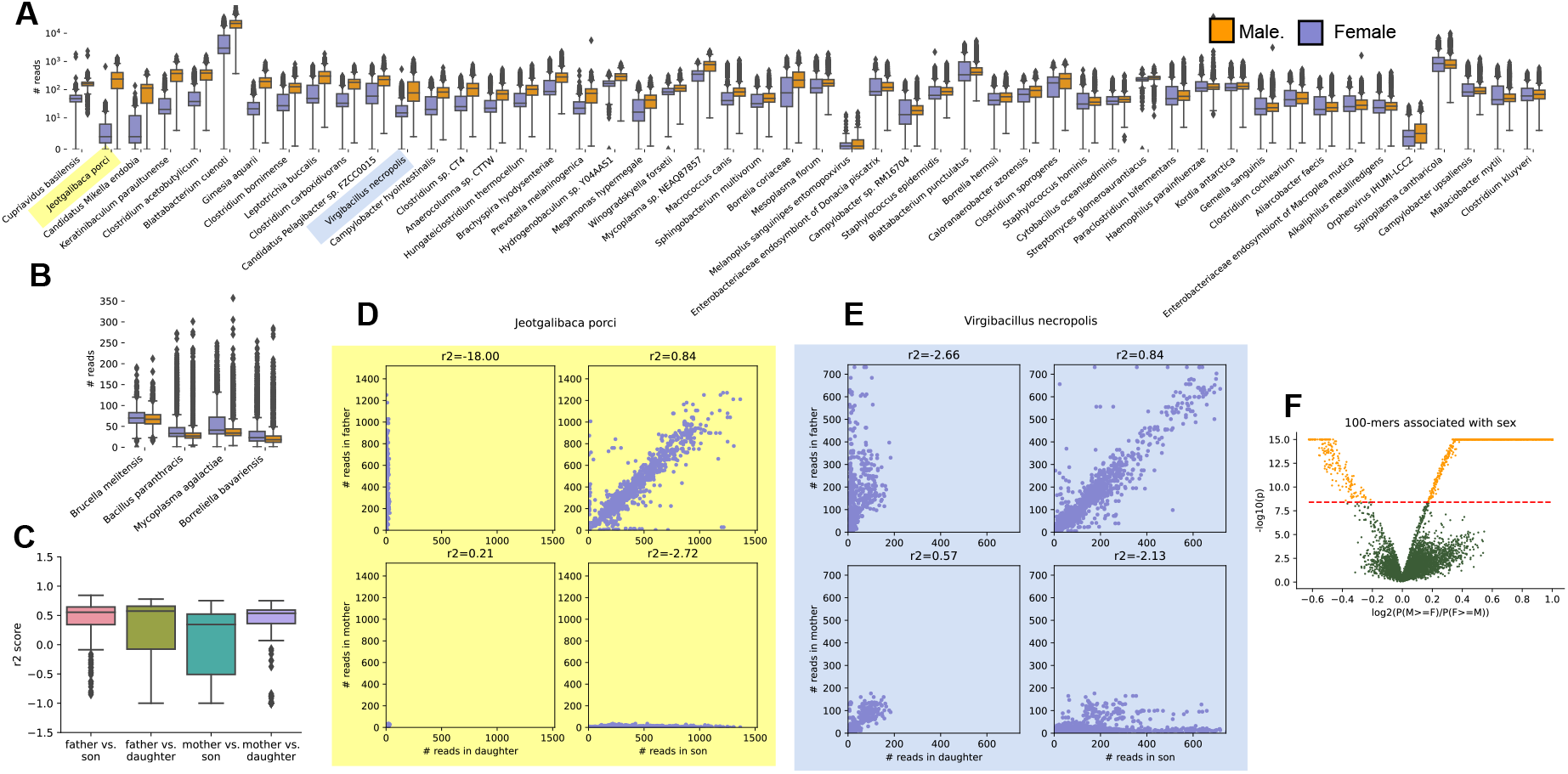
**(A)** Bacteria significantly enriched in males, subsetted to the 50 bacteria with the strongest association. Examples of bacteria with distinct inheritance patterns are highlighted in yellow and blue and shown in **D-E. (B)** Bacteria that were found to be significantly enriched in females (purple). **(C)** Boxplot of correlations between children and parents of different sexes, as measured by the *R*^2^ metric of log-normalized counts. **(D)** Example of a bacteria with inheritance patterns illustrative of y-chromosome repeats mismapping to bacterial contigs. **(E)** Example of a less clear inheritance pattern, which we hypothesize results from a sequence present both on the X and Y chromosome in varying numbers of repeats. **(F)** A sub-sampled volcano plot for the association between counts of 100-mers extracted from reads mapping to sex-associated bacteria and sex, using a paired analysis of siblings. The red line represents the Bonferonni-adjusted p-value cutoff, with statistically significant hits in orange.

Regardless of inheritance mechanism, we sought to identify particular sequences leading to these problematic mismap-pings in order to help other researchers prevent sex-biased errors in future studies. We extracted 100-basepair sequences from the reads that mapped to any of the sex-associated bacteria. We then performed a strict paired analysis using males and females of the same autism phenotype from the same family, analyzing differences in coverage of a given 100-mer. We identified 77,647 100-mers significantly enriched with males and 369 in females (adjusted p-value *<*.05). We make these k-mers publicly available and recommend that studies susceptible to such sex-biased mismappings mask reads containing these problematic sequences.

## Discussion

### Contamination Profile of iHART is similar to other NGS datasets

Many of the contaminants found in our data have been documented in other large scale genomic studies. One study [44] found several phages with abundances correlated to that of phiX, similar to what we found. The authors interpreted this as contamination of the commercial preparation of phiX174. *Mycoplasma* contamination has been found in many cell lines [45], and*Bradyrhizobium* was the most common contaminant found in WGS from the 1000 genomes project [20]. *Staphylococcus, Acinetobacter, Streptococcus*, and *Pseudomonas* have been identified as possible contaminants in several WGS studies [46, 47, 48]. Despite numerous studies cataloguing bacterial contamination in NGS, the same species seem to persist in sequencing contamination.

### Dangers of Unchecked Contamination in NGS studies and Reference Genomes

NGS of large case control cohorts become one of the most popular study designs in studies of human health. Whether hoping to identify variants in the human genome contributing to disease risk, gene expression profiles of particular phenotype, or understand causal effects of various microbiome signatures, one of the first steps in NGS pipelines is to typically to align raw reads to a set of reference databases. Unchecked bacterial contamination in NGS can compromise NGS in a variety of ways: In metagenomics studies, bacterial contamination from laboratory reagents can distort abundance counts of microbes in the samples [15], and in the worst case can lead to spurious associations between disease and microbial signature[49]. In human genetics studies, contamination mismapped to the human reference genome can lead to false variant calls [17], and different amounts of contamination across samples makes it difficult to maintain consistent coverage levels across samples. Decontamination software packages may help with some of these issues but special care must be taken to sequence a control sample for all combinations of sequencing plates and sample storage and prep, as these different experimental parameters have clear differences in bacterial contamination signatures. Meticulous paired study designs controlling for the potential for contamination in different steps of the pipeline (ie sequencing each case/control pair on the same plate, extracting and prepping the samples on the same timeline) may also reduce the risk of contamination causing a false association between microbial signature and disease. Regardless of study design, in studies of microbiota with low bacterial load, contamination from laboratory regents can limit identification of related low abundance microbes [21]. More needs to be done to understand and mitigate laboratory and reagent sources of contamination.

Poor quality reference genomes pose an additional set of risks for next gen sequencing studies. Misconstructed reference genomes that are actually chimeras of several organisms can result in incorrectly identifying which microbes are present [] and [] Incomplete reference genomes, such as the human Y chromosome, may result in mismappings from reads coming from poorly catalogued sections of genome to satisfactorily similar sequences on well-characterized reference genomes.. We have identified 10,000 100-bp sequences likely originating from the sex chromosomes that mismap to bacterial contigs. It is possible these mismappings are due to poorly constructed bacterial reference genomes that actually contain human DNA sequences [34], mismappings from Y chromosome reads as a result of its incomplete reference, or a combination of both. Regardless, we have made these problematic sequences available in a fasta file format. Many read masking and trimming tools, such as `BBTools` [50, 51] or `Trimmomatic` [52], and `Cutadapt` [53], can take in a `fasta` file of adaptor or contaminant sequences and remove reads that contain any of the problematic sequences. We recommend metagenomics or other studies performing alignment of reads derived from a human host remove reads with these problematic sequences, in order to reduce potential sex-related artifacts. This is particularly important in studies of microbiota with low bacteria-to-host-DNA ratios, such as the blood microbiome.

### Unmapped Read Space as an Untapped Data Source

Unmapped reads can constitute up to 30% of WGS data, and usually are thrown out in downstream analysis. With the wealth of WGS data that has been and continues to be generated, this unmapped read space composes several petabytes of data. We, and others [47] have shown that the unmapped read space is a valuable resource for quantifying contamination that might pollute NGS studies. The unmapped read space may also be a valuable resource for better understanding the virome [44, 54] as well as host genetic diversity [54, 55, 56], especially with the help of a well-characterized contamination profile.

## Conclusions

The unmapped read space of WGS contains information on common contaminants of WGS. Contamination profiles depend on primarily cell source type and sequencing plate. Additionally, many sequences from the Y-chromosome mismap to bacterial contigs, creating problematic sex-biased bacteria counts. The unmapped read space is a valuable resource for better understanding ubiquitous contamination patterns in WGS.

## Methods

### Extracting Unmapped and Poorly Aligned Reads

We obtained Whole Genome Sequencing (WGS) data from the Hartwell Autism Research and Technology Initiative (iHART) database, which includes 4, 842 individuals from 1, 050 multiplex families in the Autism Genetic Resource Exchange (AGRE) program 1C.

All WGS data from the iHART database have been previously processed using a standard bioinformatics pipeline which follows GATK’s best practices workflows. Raw reads were aligned to the human reference genome build 38 (GRCh38_full_analysis_set_plus_decoy_hla.fa).

We excluded secondary alignments, supplementary alignments, and PCR duplicates from downstream analyses. We extracted reads from the iHART genomes that were unmapped or mapped with low confidence. Low-confidence reads were defined as reads marked as improperly paired and with an alignment score below 100. We used alignment score rather than mapping quality in order to select for reads were likely not true alignments to the human reference genome, rather than for reads that had ambiguous alignments to hg38.

### Re-alignment

We used Kraken2 [57] to align the unmapped and poorly aligned reads to a the Kraken default (RefSeq) databases of archaeal, bacterial, human (GRCh38.p13), and viral sequences [58]. These references databases were accessed on Feb 16, 2021. Kraken2 was run on the unmapped and poorly mapped reads from each sample, using the default parameters. Because Kraken was able to map the majority of reads down to the species or strain level, Kraken classifications were aggregated by species before downstream analysis.

### F-Regression

To analyze the effect of various demographic (such as household, autism status, and sex) and experimental parameters (such as sequencing plate and sample type) on microbial and viral profile, we performed an F-regression analysis. We chose an F-regression because many variables were highly collinear with each other: for example, samples from the same household were nearly always sequenced on the same sequencing plate, autism is much more prevalent in males, and the same sample types were normally collected from households. For each microbe, we built an ordinary least squares (OLS) model, using as our regressor an indicator matrix of sample type, sex, child vs. parent, autism status, sequencing plate, household/family, and sample id, and as our response variable the log-normalized counts of microbes (with pseudo-counts of 1). Using the `statsmodels` library, [59] we then ran a forward OLS regression in which we iteratively selected the regressor features that best explained the previous model’s residuals, and ceased adding features when the ANOVA score between the previous and new models was no longer statistically significant (p*<*.05). [60]

### Y-Chromosome Mismapping Analysis

Using the F-regression, we found that many microbes were significantly associated with sex (162 species were enriched in males and 4 species were enriched in females. Hypothesizing that such mismappings were due to mismappings of repetitive regions on the X or Y chromosome, we analyzed inheritance patterns, looking at the correlation between children and parents using the r2 score (as shown in Fig. 4. Furthermore, we sought to identify specific subsequences that cause these problematic bacterial classifications. From the collection of reads that aligned to the bacterial reference contigs associated with sex, we extracted and counted the occurrence of 100-basepair k-mers in every sample. We counted the 100mers using the highly parallel k-mer counter `jellyfish`. We chose 100-basepair k-mers because assuming a uniform distribution of 150b reads across the human genome at 30x coverage and a trivial sequencing error rate, 100 bases is the longest length of a k-mer with over a 99.5% chance of being captured within the 150 bases of at least one read in an individual. To reduce k-mers generated by sequencing error or low frequency genetic variants, we filtered to 100-mers that occurred at least twice in at least two samples. In order to test the null hypothesis that these subsequences show equal occurrences in males and females, we then performed a paired test between males and females siblings within the same family with the same autism status (to stringently weed out ancestry and disease phenotype as confounding variables). We reported the 100-mers that had a Bonferonni-adjusted p-value *<*.05, and make them publicly available for access in a “‘fasta”‘ format that can easily be access by read trimming and masking tools. These sequences are available at the link described below.

## Availability of data and materials

The iHART dataset is available upon reasonable request at http://www.ihart.org/home.

The complete list of reference genomes used for Kraken realignment can be found at http://github.com/briannachrisma

The sequences associated with sex can be found at http://github.com/briannachrisman/blood microbiome/public and http://github.com/briannachrisman/blood_microbiome/public_data/y_sequences.fasta

## Consent for publication

All authors reviewed this manuscript and gave consent for publication.

## Authors’ contributions

BSC wrote the manuscript. BSC, CH, and JJ wrote and developed the source code. DPW, NS, KP, and PW contributed to the conception and design of the study, facilitated collection and sequencing of autism dataset, and participated in the analysis of the results. All authors read and approved the final manuscript.

